# 3D co-culture of macrophages and fibroblasts in a sessile drop chip for unveiling the role of macrophages in skin wound-healing

**DOI:** 10.1101/2023.01.04.522690

**Authors:** Xiaoyan Lyu, Feiyun Cui, Hang Zhou, Bo Cao, Minghui Cai, Shulong Yang, Bangyong Sun, Gang Li

**Author notes:** Correspondence: Xiaoyan Lyu, Gang Li. These authors contributed equally to this work (Xiaoyan Lyu, Feiyun Cui, Hang Zhou).

## Abstract

Three-dimension (3D) cell co-cultural spheroids exhibit enhanced cellular functions and they can mirror in-vivo microenvironments. Herein, a sessile drop chip was developed to construct 3D spheroids for mirroring the wound healing microenvironment. The sessile drop chip holds the superhydrophobic surface of each microwell which can facilitate cell suspensions transfer to spheroids through the offset of surface tension and gravity, and each microwell has a cylinder hole that offers adequate oxygen to spheroids. It was demonstrated that the 3T3 fibroblast spheroid and the 3T3 fibroblast/M2-type macrophage co-culture spheroid can be formed and remained the physiological activity within nine days. 3D morphology of spheroids was reconstructed using the transparent processing technology and Z-stack function of confocal microscopy. Characteristics of proliferation and differentiation were analyzed by using nano antibody-based 3D immunostaining assay. Results revealed that M2-type macrophages can promote the proliferation and differentiation of the 3T3 fibroblast spheroid. This study presented a novel affordable platform for developing 3D spheroids and provides a 3D model for investigating the macrophages-associated wound healing microenvironment.

## 1. Introduction

Macrophages and fibroblasts play critical roles in the wound healing process ^1^. When skin injury occurred, pro-inflammatory macrophages, which are traditionally referred to as “M1” macrophages, were employed to clean bacteria, foreign debris, and dead cells in the wound ^2^. After the inflammatory phase, the predominant macrophage population was changed to the wound healing phenotype called “M2” macrophage. It yields numerous growth factors and stimulates tissue fibroblasts to differentiate into myofibroblasts, which could facilitate wound contraction and closure ^3^. After that, the dominant M2 macrophage populations exhibited a mostly anti-inflammatory phenotype ^4^, likewise, fibroblasts produce ECM molecules that regulate tissue strength and resilience ^5^. The research of these biological mechanisms is of great significance for guiding the treatment of wounds. Although considerable advances have been achieved, the research on the interrelationship between macrophages and fibroblasts still remains a challenge for further exploring wound-healing mechanisms and related drug development.

Traditionally, trans-well migration assay and scratch assay are commonly used bioassays to observe and study interactions between cells ^6^. Both of them are low throughput and the types of cells that can be observed at the same time are limited ^7,8^. Therefore, these methods are not competent for observing the regulatory effect of macrophages with different polarities on a variety of skin intrinsic cells. The development of alternative platforms for studying the interrelationship between these two cells is expected.

Most cells in the physiological environment in-vivo are continuously exposed to physical and chemical interactions with neighboring cells in a three-dimension (3D) space. This distinctly regulates the intracellular responses and extracellular interactions which are difficult to recreate in a two-dimension (2D) monolayered cell culture platform. In a common 2D environment, cells have more surface area in contact with the plastic and culture media than with other cells forcing them into a polarization that does not reflect physiological conditions in-vivo ^9^. Meanwhile, most cell interaction analyses are performed in-vitro 2D cultures, which disregard the complexity of interactions that occur in-vivo ^10^. Indeed, 2D cultured primary human hepatocytes (PHHs) show rapid declines in critical phenotypic functions ^11^. In contrast, cells cultured in 3D multilayer conditions grew in all directions and interacted with neighboring cells through biochemical and mechanical cues ^12, 13^ and can be manipulated to mimic cell interactions in-vivo ^14^. Hence, 3D cell culture or cell co-culture is an alternative technology for cell interaction study ^15^, cellular morphology ^16^, functions ^17^, overall behavior^18^, as well as molecular biology ^19^.

Usually, 3D culture techniques can be categorized into scaffold-based and scaffold-free types, as well as hybrid 3D culture models in which formed spheroids were incorporated into a 3D polymeric scaffold ^20–22^. Scaffold-based culture technologies provide physical support, ranging from simple mechanical structures to extracellular matrix (ECM)-like matrices, on which cells can aggregate, proliferate, and migrate. They include microfluidic devices, polymeric hard scaffolds, hydrogel scaffolds, microplates, micropatterned surfaces, and spheroids. Scaffold-free techniques rely on the self-aggregation of cells in specialized culture plates, such as hanging drop microplates ^23^, low adhesion plates with ultra-low attachment coating that promotes the spheroid formation ^24, 25^, and micropatterned plates that allow for microfluidic cell culture ^26^. The development of spheroids is the most popular and convenient technique for 3D cell co-culture. Spheroids, especially cell co-culture spheroids can recapitulate the physiological characteristics of tissues or tumors concerning cell-cell contact, and can produce their own ECM, allowing for the natural cell-matrix interactions ^27^.

Herein, a novel cost-effective sessile drop chip was proposed to investigate the macrophage-modulated fibroblasts spheroid in the skin repair microenvironment. As shown in Fig. 1, By modifying a recessed surface of the sessile drop chip by one a superhydrophobic solution, cell suspensions can be entered into an array of superhydrophobic microwells and stabilized under the combined action of surface tension and gravity. With a thru-hole at the bottom of each well, the cell spheroids can acquire adequate oxygen to avoid apoptosis. The powerful tool was applied for spheroid culture and analysis. M2 macrophages were commixed with fibroblast in the sessile drop chip to better understand the effects of macrophages on fibroblast in the wound healing process.

**Figure 1.**
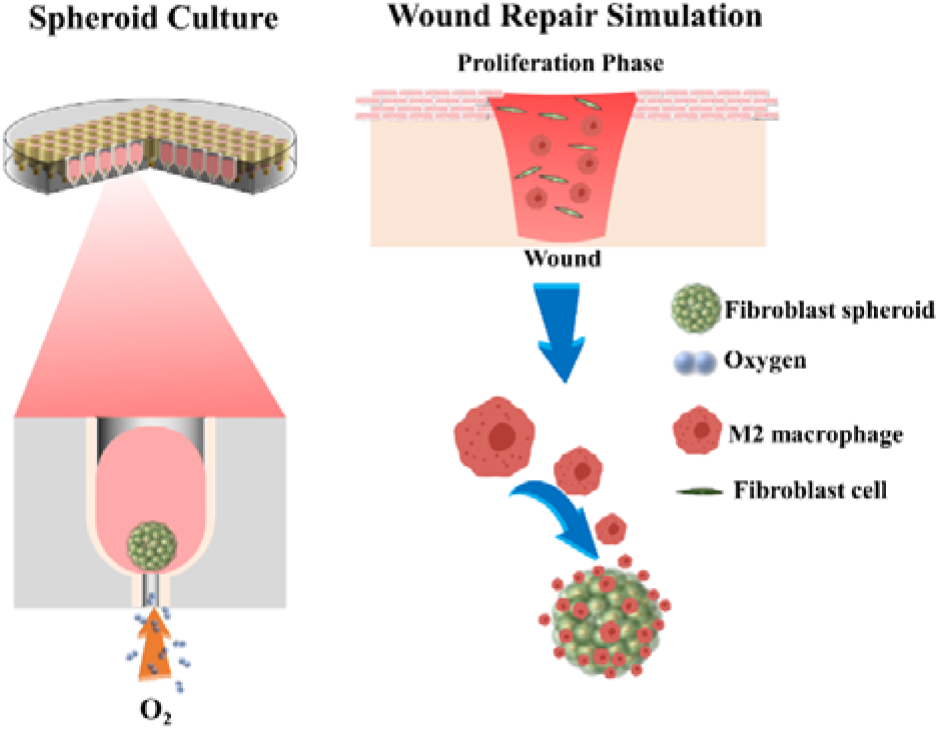
Schematic diagram of the proposed sessile drop chip, single well with oxygen hole, and fibroblast and macrophage associated wound repair pattern.

## 2. Results and Discussions

### 2.1 Construction of the 3T3 fibroblast spheroid in the superhydrophobic sessile drop chip

Spheroids can be grown to a size where they display oxygen and nutrient gradients similar to tissue. The sessile drop chip was fabricated by computer numerical control (CNC) milling with polymethyl methacrylate (PMMA) plate and coated with octadecyl trichlorosilane (OTS) solution to form a superhydrophobic surface (Fig. S1). There are two advanced features in the sessile drop chip: (1) The superhydrophobic surface of each microwell can facilitate cell suspensions transfer to spheroid through the offset of surface tension and gravity and (2) A cylinder hole at the bottom of each microwell, which can offer adequate oxygen to spheroids.

Firstly, the sessile drop chip was applied for culturing of 3T3 fibroblast spheroids to verify its function and explore optimization experimental conditions. To obtain high-quality spheroids, a number of cells was dripped into a 10×10 microwell of the sessile drop chip (Fig. 2A). Due to the presence of superhydrophobic coating, the cells cannot attach to the surface of the microwell, thus spontaneously formed a spheroid in the bottom of each microwell (Fig. 2B). The size, morphology and biological activity of spheroids depend on the initial number of cells seeded. To confirm the optimal quantity of 3T3 cells for 3D cultivation, 5000 to 80000 cells/mL were added into a single well, respectively. As shown in Fig. S2, the cells in a single microwell formed several non-uniform size spheroids when adding 5000 cells/mL, 10000 cells/mL, and 80000 cells/mL. They are unfeasible for further study. To obtain a single easy-measured spheroid, 20000 and 40000 cells/mL were suitable. The amount of 40000 cells/mL was selected because the spheroid is larger and can be measured conveniently in further experiments. To investigate the morphology of 3T3 fibroblast spheroid, fluorescent microscopy was used to measure the area change at the XY axis. Interestingly, the 3T3 fibroblast spheroid was contracted when cultured from day 1 to day 5 in 2D view, and then it subsisted in a stable condition (Fig. 2c). Fig. 2d suggested that the average area of 3T3 fibroblast spheroid was decreased from 148000 μm^2^ to 75000 μm^2^ from the contraction stage of day 1 to day 5, after that, the average area maintained at almost 75000 μm^2^.

**Fig 2.**
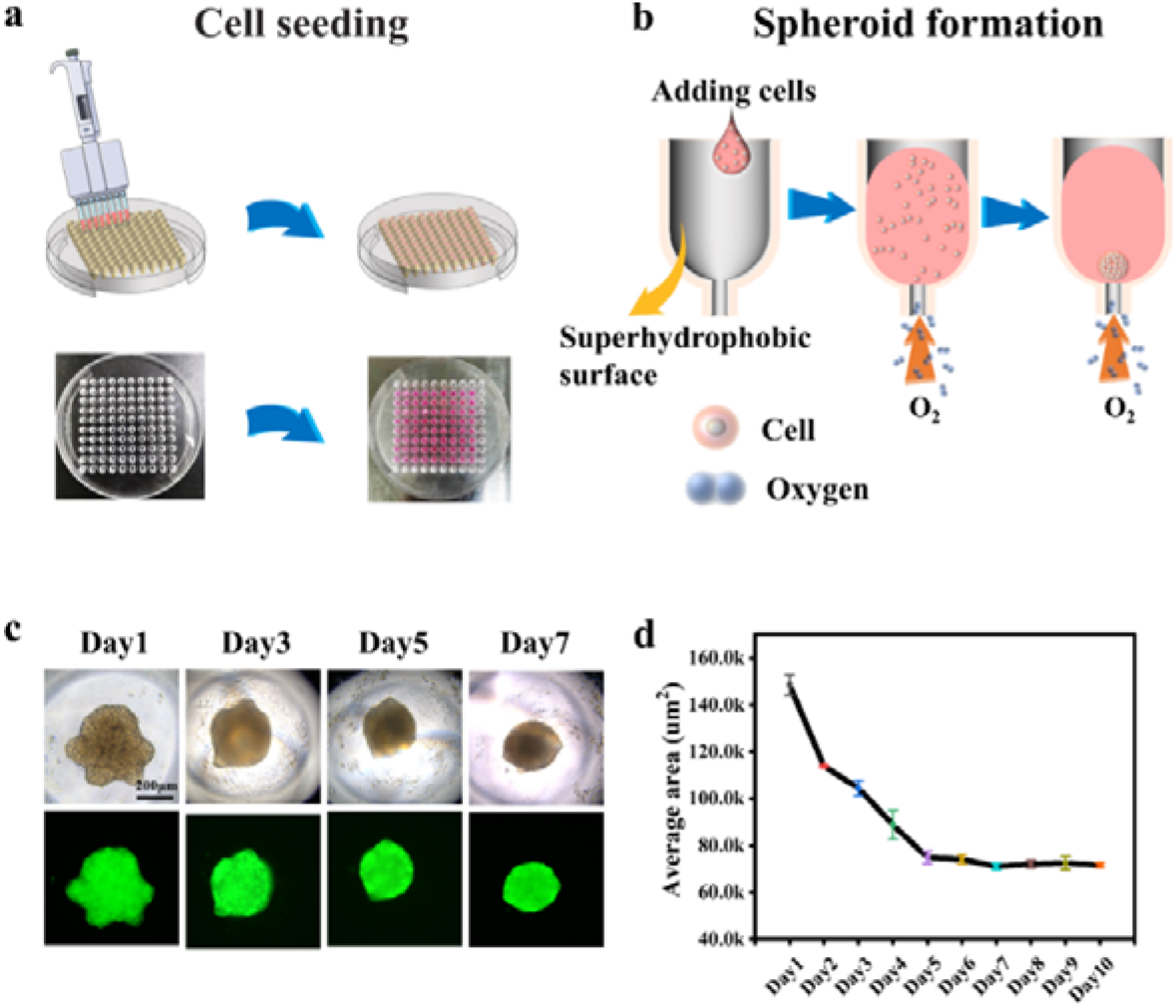
Construction of 3T3 fibroblast spheroid in the sessile drop chip and optimization of culture condition. (a) Schematic Diagram of seeding 3T3 cells to a sessile drop chip. (b) A 3T3 fibroblast spheroid formed spontaneously in a microwell. (c) The microscopic images of 2D areas of a 3T3 fibroblast spheroid from day 1 to day 7, (d) the 2D average areas chart of 3T3 fibroblast spheroid from day 1 to day 10.

### 2.2 3D reconstruction of 3T3 fibroblast spheroid

According to the 2D image analysis, the 3T3 fibroblast spheroids were constricted and then sustained a stable area. It’s not a common phenomenon compared with reported tumor spheroids, in which the diameter in the 2D view will increase within a reasonable time range ^28–30^. Hence, it was needed to further characterize the organoid in a 3D view. As shown in Fig. 3a, a transparent spheroid was obtained after washing it with clearing reagents. It was demonstrated that the treated spheroid was limpid in the bright field, whilst the fluorescence intensity was not disturbed or affected compared with the non-treated group (Fig. S3a). Followed obtaining transparent spheroid, the Z-stack function of confocal microscopy was utilized to reconstruct the 3D morphology of the 3T3 fibroblast spheroid (Fig. 3a). As shown in Fig. 3c and 3d, 3D models of 3T3 fibroblast spheroid from day 1 to day 9 were constructed with Z-stack. In Fig. 3c, it can be observed that the altitude of the Z axis on day 1 was higher. The orthogonal view (Fig. 3d) revealed that 3T3 cells were not shaped as a spheroid on day 1 and there was a concave area at the center of the cell clumps. Then the concave area gradually disappeared within 3 days, and the area of the XY axis dwindled simultaneously from day1 to day 5, the similar constructions were also observed in the XZ view and YZ view. Then the 3T3 fibroblast spheroid ceases contraction (Fig. 3d 3T3 day 5-day 7) in three views (XY, XZ, YZ) and the spheroid volume rarely changes in period of day 5 to day 9. Fig. 3b showed the whole spheroid volume change, the volume decreased from 8.3 mm^3^ to 2.6 mm^3^ from day1 to day5, then kept stable at 2.6 mm^3^ from day 5 to day 9. These changes suggested that the contraction process of a 3T3 fibroblast spheroid existed from day1 to day 5 and a steady period existed from day 5 to day 9. The panoramic variations of the 3T3 fibroblast spheroid were presented in Fig. S4 and Fig. S5.

**Fig. 3.**
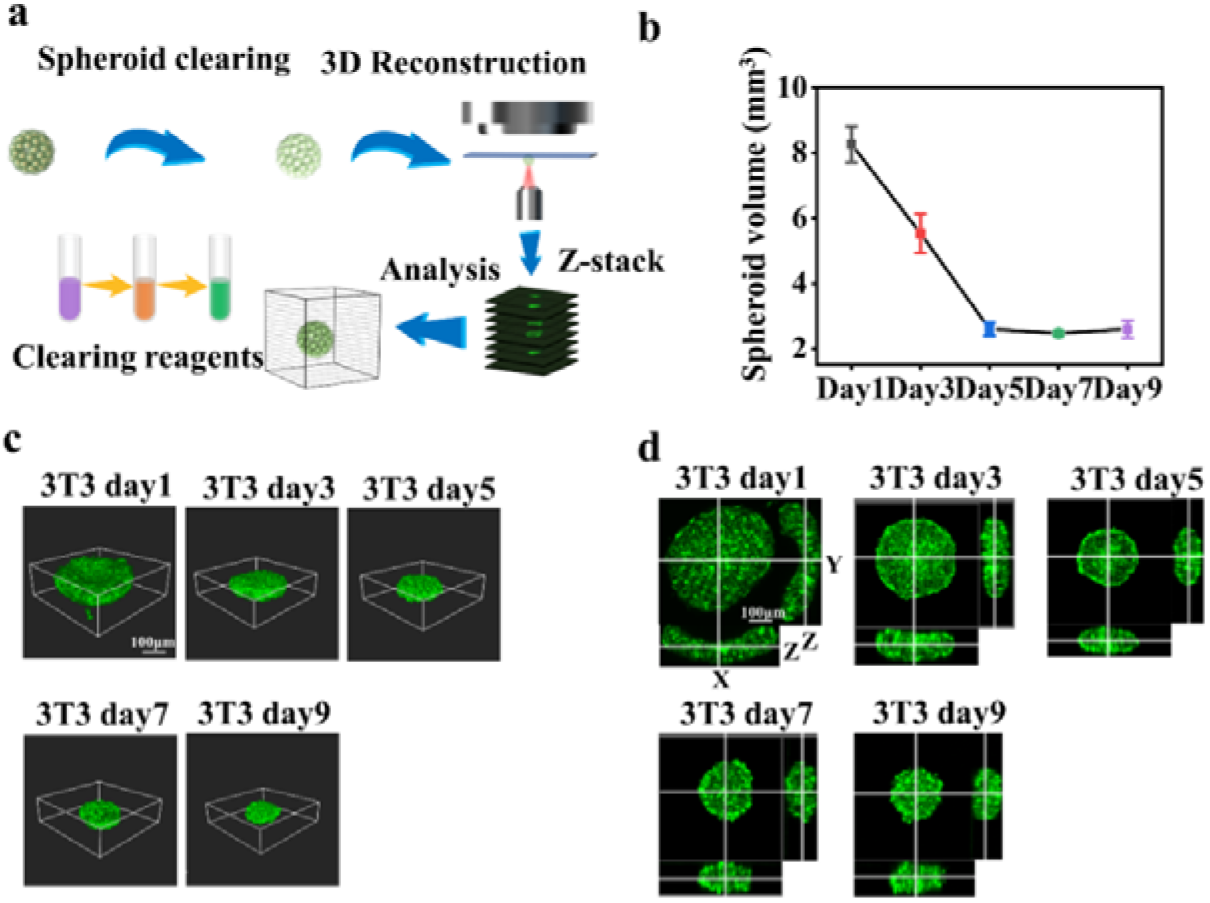
3D reconstruction of 3T3 fibroblast spheroid from day1 to day9. (a) Scheme of spheroid clearing and 3D reconstruction. (b) The volume change of the spheroid from day 1 to day 9. (c) 3D images of 3T3 fibroblast spheroid from day 1 to day 9. (d) Orthogonal images of spheroid from day 1 to day 9.

### 2.3 Proliferation and differentiation of 3T3 fibroblast spheroid

The proliferation and differentiation of 3T3 fibroblast spheroid from day1 to day 9 were investigated by immunofluorescent assay (IFA). It is quite different to perform the immunostaining assay in a 3D spheroid compared with a 2D cell. The thickness of the 3D spheroid in the Z axis was large and the antibody (especially the secondary antibody) cannot easily diffuse in the interior of the spheroid. As shown in Fig. S6, the immunostaining condition was optimized by using the nano-secondary antibody which is 10 times smaller than the normal secondary antibody. The whole spheroid was completely stained with Ki67 when using nano-secondary antibody compared with normal secondary antibody. Hence, it is applicable to investigate the proliferation and differentiation ability of the 3T3 fibroblast spheroid using 3D immunostaining. As shown in Fig. 4a and Fig. S7, 3T3 fibroblast spheroids from day 1 to day 9 were stained with Ki67 and tubulin, respectively. On day 1, the fluorescence of Ki67 and tubulin were distributed in the whole spheroid. On day 5, it was observed that Ki67 was dimmed in the internal area of the spheroid which indicated that the proliferation ability of the inner area was decreased. From day 7 to day 9 (Fig. 4a, S7), the fluorescence of Ki67 was revived evidently and the Ki67 are at the verge of these spheroids was broader and the cells in this area showed a growing trend compared with day 5. Meanwhile, the fluorescence of tubulin was changed slightly. Therefore, it was supposed that the proliferation ability had gradually waned in the interior of the 3T3 fibroblast spheroid in the contracting phase (day 1-day 5), then it was slowly recovered because the cells located at the exterior of the spheroid started proliferation in the stability phase (day 5-day 9). The average fluorescence of Ki67 and tubulin was measured respectively and the fluorescence intensity ratio of Ki67 and Tubulin was calculated (Fig. 4b). It showed that the ratio declined from 0.98 to 0.72 from day 1 to day 5, then recovered to 0.83 from day 5 to day 9. All immunostaining results of Ki67 and tubulin on day 1-day 9 were presented in Fig. S7.

**Fig 4.**
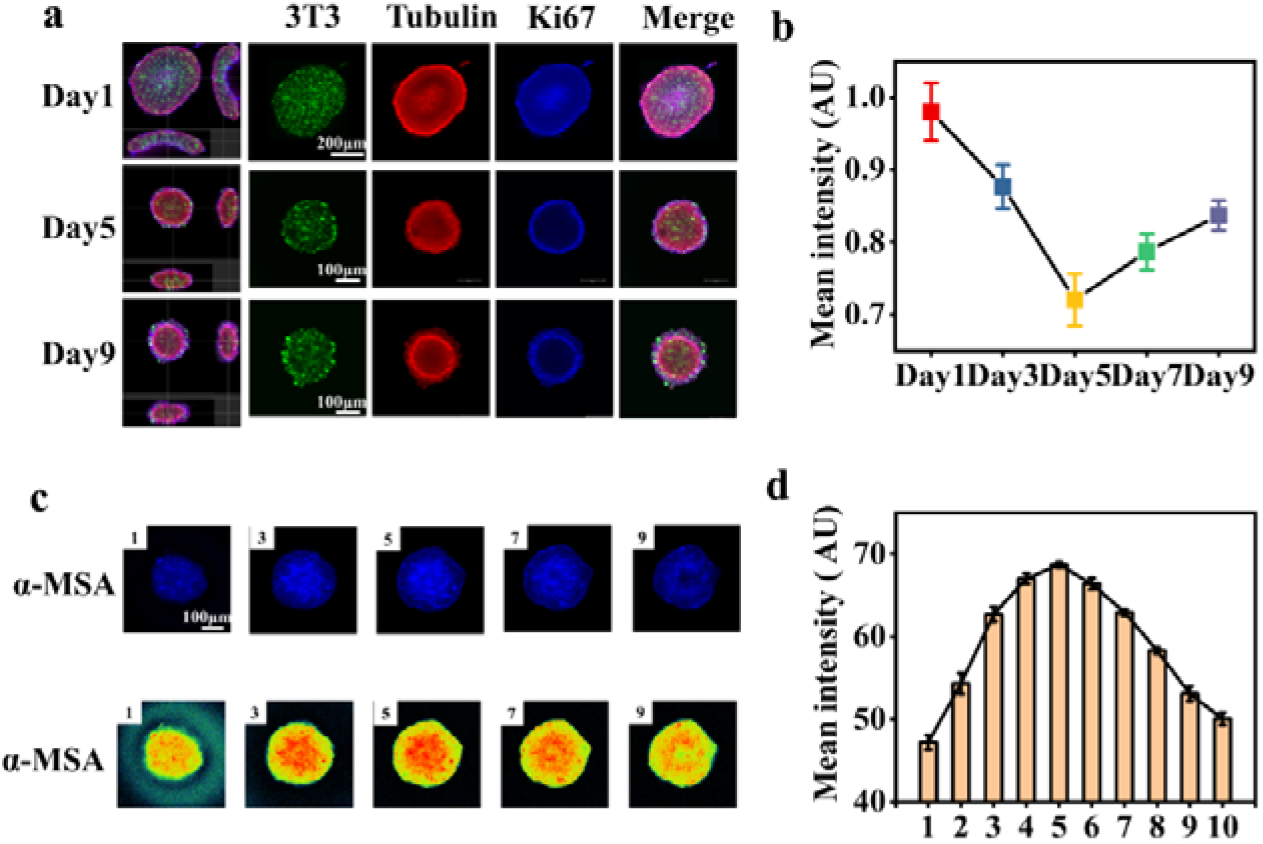
Proliferation and differentiation of 3T3 fibroblast spheroid using 3D IFA. (a) Immunostaining of Ki67 and Tubulin in 3T3 fibroblast spheroid on day 1, day 5, and day 9. (b) The ratio of the average intensity of Ki67/Tubulin on day1, 3, 5, 7, 9. (c) Fluorescence images and heatmaps of five sequential sections of α-SMA expression in the 3T3 fibroblast spheroid. (d) Average fluorescence intensity of 10 sections (the images of 10 sections were presented in Fig. S8).

Usually, 3T3 fibroblasts tends to differentiate into myofibroblasts in the wound healing microenvironment ^31^. Herein, the cell differentiation ability of the 3T3 fibroblast in the 3D spheroid was demonstrated using a differentiation marker α-smooth muscle actin (α-SMA). In Fig. S8, 10 sequential sections of a 3T3 fibroblast spheroid on day 4 were emphasized (only showed 5 sections, the whole sections were displayed in Fig. 4c). They were arranged from one side of the spheroid to another side of the spheroid on the Z-axis. It was found that the fluorescence of α-SMA in the fifth image (which is the intermediate section of the spheroid) was brighter than the peripheral area. It reveals that the 3T3 fibroblast were differentiated well in the center of the spheroid. Furthermore, the expression of α-SMA in the middle section is stronger than that in the edge section. The result is more obvious in heatmaps (Fig. 4c). In Fig. 4d, the average value of fluorescence intensity of α-SMA in all 10 sequential sections was calculated. It showed a tendency that the fluorescence was higher in the middle and lower on both sides. Furthermore, it proved that the central 3T3 fibroblast were differentiated well. In addition, the fluorescence intensity of α-SMA from day 3 to day 6 was evaluated and presented in Fig. S9. It was demonstrated that the cells which expressed higher α-SMA were widely distributed in the entire area of the 3T3 fibroblast spheroid on day 3 and gradually scattered at the center on day 4 to day 6.

### 2.4 Co-culture spheroid of 3T3 fibroblast and M2-type macrophage for simulate the wound healing microenvironment

Fibroblasts and macrophages are two critical cells in wound healing ^32, 33^. Studying the interrelationship between these two cells based on the novel developed platforms is crucial for the exploration of wound-healing mechanisms and related drug development ^34, 35^. Herein, the M2-type macrophage (RAW 264.7) and 3T3 fibroblast were co-cultured. The M2-type macrophage expressed the red fluorescence protein (RFP) and 3T3 expressed the green fluorescence protein (GFP) were employed in this study. According to our laboratory’s studies ^36, 37^, the RAW 264.7 M2-type macrophage can subsist for 5-6 days after stimulating with IL-4. Thus, the co-culture system was recorded within 5 days. As shown in Fig. 5b, when culturing on day 1, the clumps of 3T3+M2 co-culture spheroids were thinner compared with 3T3 fibroblast spheroids (Fig. 5a). Both of them showed a contracting trend with the passage of time. Furthermore, the M2-type macrophage gradually wrapped the 3T3 fibroblast spheroid especially on day 3-5 (Fig. 5c).

**Fig 5.**
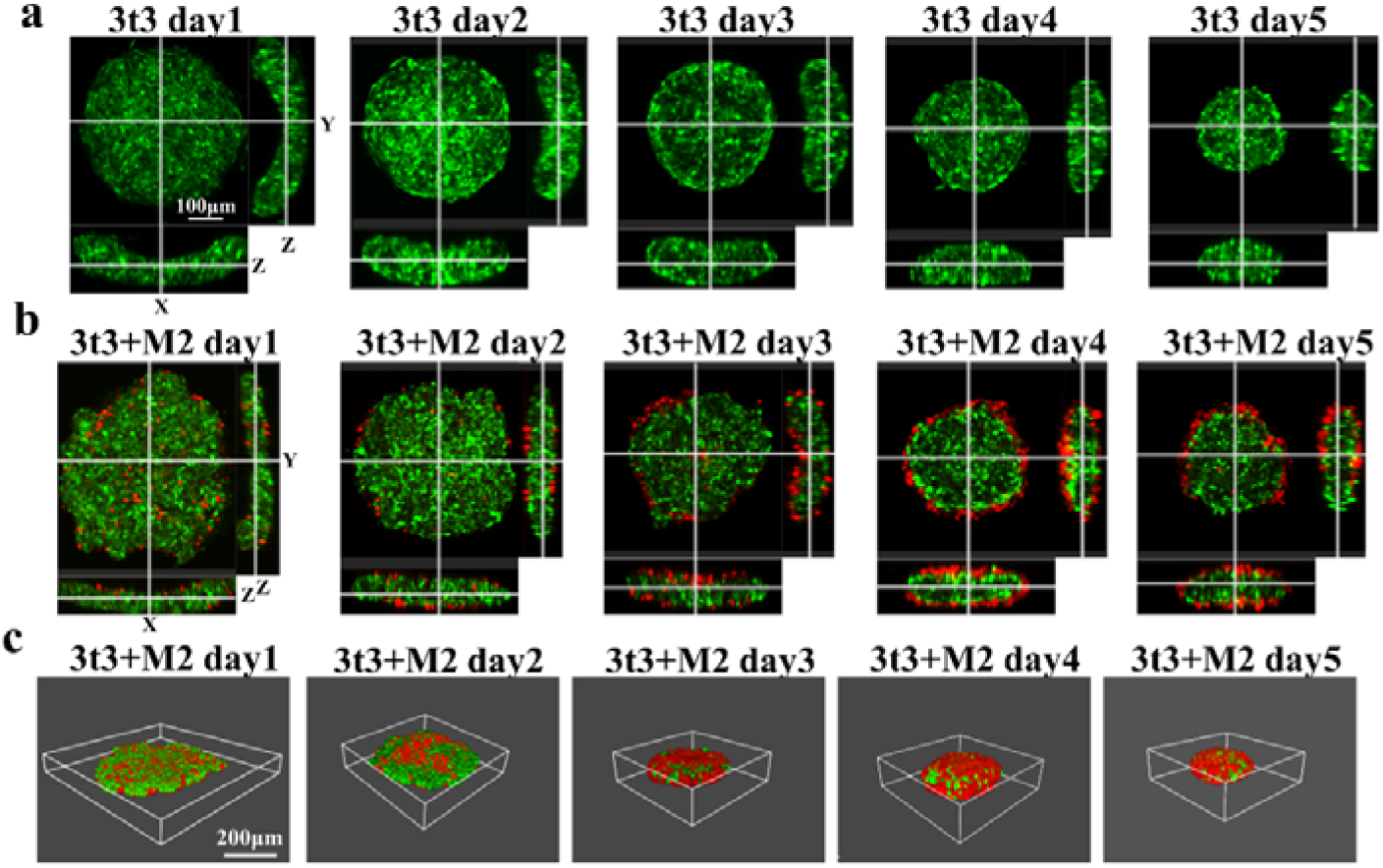
Co-culture of M2-type macrophage (RAW264.7 cell) and NIH-3T3 fibroblast cell. (a) Orthogonal images of 3T3 fibroblast spheroids from day 1 to day 5. (b) Orthogonal images of 3T3+M2 co-culture spheroids from day 1 to day 5. (c) 3D images of 3T3+M2 co-culture spheroids from day 1 to day 5.

### 2.5 Proliferation and differentiation of fibroblasts in co-cultured spheroid

To investigate the effects of M2-type macrophage on the fibroblast 3T3 in spheroid conditions, the proliferation ability of 3T3 in the 3T3+M2 co-culture spheroid was studied. Fig. S10 showed that the 3T3+M2 co-culture spheroids have a consistent tendency compared with 3T3 fibroblast spheroids. The Ki67 fluorescence intensity of both spheroids declined from day 1 to day 5. It suggested that the proliferation ability of 3T3 fibroblast spheroids was not improved when co-culturing with M2-type macrophage. The differentiation of 3T3 day 4 and 3T3+M2 day 4 were evaluated (Fig. 6a). The sections of 3T3 and 3T3+M2 were selected at the same position on the Z axis. Then the fluorescence of α-SMA of the above-mentioned spheroids was measured using the same excitation light and absorbed light. The result (Fig. 6b) showed the 3T3+M2 was brighter than 3T3, which means the 3T3 cells differentiated better when co-cultured with the M2-type macrophage. The average intensity of α-SMA in the 3T3 fibroblast spheroids and 3T3+M2 co-culture spheroids were calculated. Fig. 6c suggested the intensity of the 3T3+M2 co-culture spheroids (67) is higher than the 3T3 fibroblast spheroids (44) with P<0.01.

**Fig 6.**
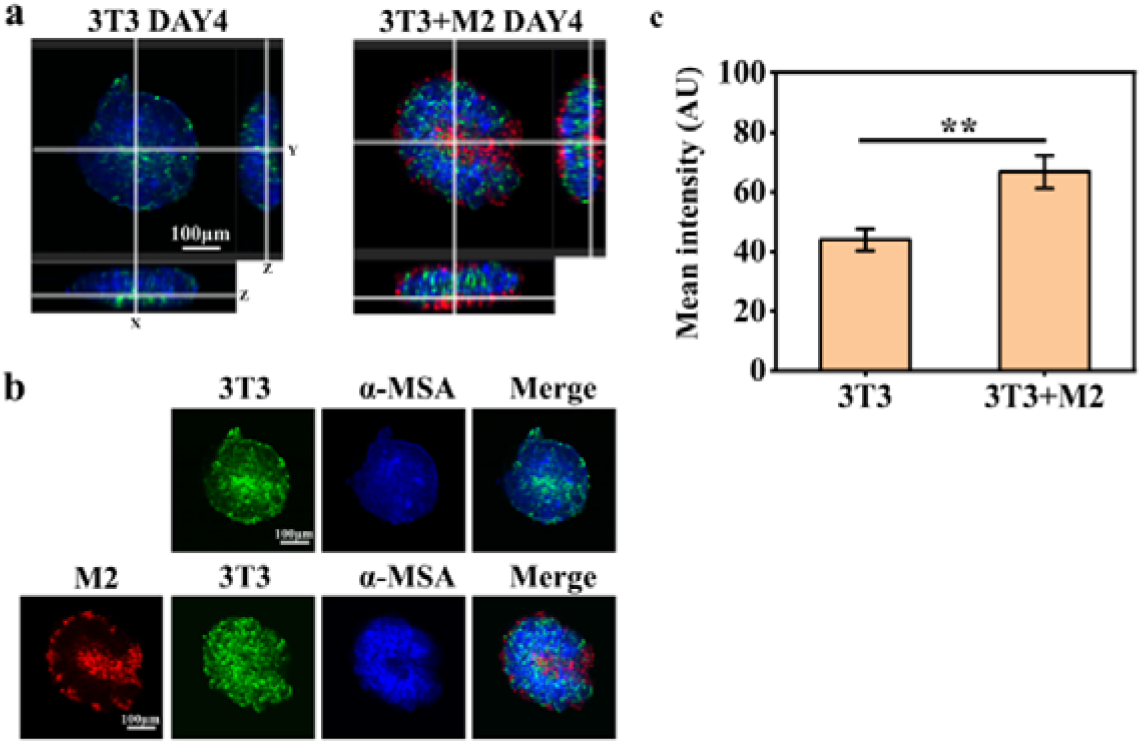
Comparison of proliferation and differentiation of 3T3 fibroblast spheroids and 3T3+M2 co-culture spheroids using 3D immunostaining. a) Orthogonal images of 3T3 fibroblast spheroids and 3T3+M2 co-culture spheroids on day 4. b) Immunostaining images of α-SMA of 3T3 fibroblast spheroids and 3T3+M2 co-culture spheroids on day 4. c) The column chart of α-SMA intensity between 3T3 fibroblast spheroids and 3T3+M2 co-culture spheroids.

## 3 Conclusions

In the study, 3D co-culture spheroids of 3T3 fibroblast and M2-type macrophage were developed for the reconstitution of wound healing microenvironment on a low-cost and user-friendly sessile drop chip. Each microwell in the sessile drop chip holds the superhydrophobic surface and has a cylinder hole which was used to supply adequate oxygen to spheroids. The concentration of inoculated cell was optimized to 4000 cells/mL and the formed single 3T3 fibroblast spheroid and 3T3 fibroblast/M2-type macrophage co-culture spheroid can remain physiological activities within nine days. 3D morphology of spheroids was characterized by transparent processing technology and the Z-stack function of confocal microscopy. They were turn out to be in the shape of a pie rather than a perfect sphere. Proliferation and differentiation characteristics were analyzed by the nano antibody-based IFA of Ki67/Tubulin and α-SMA, respectively. It was demonstrated that M2-type macrophages can promote the proliferation and differentiation of the 3T3 fibroblast spheroid. The developed sessile drop chip and related methodologies could been easily expanded to other multicellular spheroids research and further promote the progress of 3D cell biology.

## 4. Experiments

### 4.1 Materials and reagents

Octadecyltrichlorosilane (T821379) and hexane (H811147) were ordered from Macklin. Dulbecco’s modified Eagle’s medium (DMEM), fetal bovine serum (FBS), penicillin-streptomycin, trypsin, and collagenase were obtained from ThermoFisher Scientific. Interleukin-4 (IL-4) were provided by Peprotech (Rocky Hill, CQ). A tissue clearing kit was obtained from Nuohai Life Science (China). Ki67 antibody (Cat No. 28074-1-AP), Alexa 568 nano secondary antibody (Cat No. sms1AF568-1), and Alexa 647 nano secondary antibody (Cat No. srbAF647-1) were purchased from Proteintech. Alexa 555 secondary antibody (A32732) was purchased from ThermoFisher Scientific. Tubulin antibody (HC101-02) was purchased from Transgenic. α-MSA antibody (19245S) was purchased from Cell Signaling Technology.

### 4.2 Fabrication and surface coating of the sessile drop chip

The 3D culture chip was fabricated based on the scheme described in Figure S1. The device was constructed via two elements: (1) a PMMA microplate containing repellent microwells array that holds cell droplets, and (2) a pair of pads used for gas exchange between spheroids and incubation environment. As shown in Fig. S1a, using the computer numerical control (CNC) milling technique, a 6 mm-thick polymethyl methacrylate (PMMA) plate was drilled to obtain a 10×10 array of microwells (4 mm in diameter and 5 mm in depth) with hemispherical bottoms and centered bottom thru-holes (1.5 mm in diameter). Next, the microplate was immersed in the OTS-based coating solution ^27^ to acquire a microscale or nanoscale hierarchical superhydrophobic surface (Fig. S1b). Finally, the superhydrophobic modified chip was bonded to a couple of PMMA strips using double coating tape (Fig. S1c).

### 4.3 2D cell culture

3T3-NIH fibroblast cells and RAW 264.7 macrophage cells were cultured at 37 °C and 5% CO_2_ and maintained in growth media containing Dulbecco’s Modified Eagle’s Medium (DMEM; Cat. D5648, Sigma, MO, USA) supplemented with 10% (v/v) fetal bovine serum (Cat. 10500, Gibco, Thermo Fisher Scientific, MA, USA) and 2% (v/v) penicillin-streptomycin (Cat. DE17-602E, Lonza, Basel, Switzerland). For differentiation, RAW 264.7 macrophage cells were stimulated with IL-4 (40 ng/ml) for 24 h to obtain M2 polarization.

### 4.4 3D spheroid culture

The superhydrophobic sessile drop chip was immersed in 75% alcohol and exposed to ultraviolet for 1h to sterilization. 3T3-NIH cells were washed with PBS and digested with 0.25% trypsin after growing to 80% confluence. Then cells were resuspended with medium and seeded into a superhydrophobic sessile drop chip for 3D cell culture with a gradient of cell density (5×10^3^, 1×10^4^, 2×10^4^, 4×10^4^, 8×10^4^ cells/mL per well). Thereafter, the sessile drop chip was put in a humid dish and cultured at 37 °C and 5% CO_2_ and maintained with DMEM medium supplemented with 10% FBS. The procedure of 3T3-NIH cell and RAW 264.7 M2-type co-culture were similar to the above method. Briefly, 4×10^4^ cells/mL were mixed with M2 macrophage (4×10^2^) and seeded into a single well in the plate. The images of spheroids were obtained from fluorescence confocal microscopy.

### 4.5 Clearing of the NIH-3T3 fibroblast spheroid and NIH-3T3 and RAW264.7 Co-culture spheroid

After being grown for 7 days, the spheroids were collected and fixed with 4% paraformaldehyde for 4 h, then cleared with a tissue clearing kit. According to the manufacture protocol and our optimization fixed spheroids were added into 1 mL rapid decolorization and degreasing reagent for 12 hours, then sent to 1 mL decolorization and degreasing reagent for 12 hours. Finally, spheroids were mixed with a 200 μL clearing reagent and observed the transparency via confocal microscopy. The spheroids were preserved in 1X PBST for further immunofluorescence staining.

### 4.6 3D immunostaining of NIH-3T3 fibroblast spheroid and NIH-3T3 and RAW264.7 Co-culture spheroid

After the spheroid clearing procedure, NIH-3T3 fibroblast spheroids were immunostaining based on immunofluorescence protocol. Firstly, the spheroids were washed with 1X PBST 3 times, then blocked with 10% BSA overnight at 4 □. Then adding washed spheroids to diluted 400 μL primary antibody solution (Ki67, Tubulin, α-MSA were diluted at a ratio of 1:100) and incubated overnight at 4 □, thereafter, washing the spheroids three times with PBST and adding nano-second antibody into spheroids for 6h at 4 □. Finally, spheroids were observed with confocal microscopy and reconstructed 3D model with Z-stack function. The immunostaining and observation of NIH-3T3 and RAW264.7 co-culture spheroid were consistent with the above methods.

### 4.6 Data analysis

All images were acquired from Leica Las X and analyzed with ImarisViewer and FIJI. All data were expressed as mean□±□standard deviations (SD). The student’s t-test was used to statistically analyze the data obtained from the experiments. Significant differences are indicated as *, p□<□0.05; **, p□<□0.01; ***, p□<□0.001.

## Acknowledgments

This work was supported by the National Natural Science Foundation of China (81871567) by Prof. Xiaoyan Lyu.

## Notes

### Competing Interest Statement

The authors have declared no competing interest.

