## Supplemental Material for "3D co-culture of macrophages and fibroblasts in a sessile drop chip for unveiling the role of macrophages in skin wound-healing"

* Correspondence:

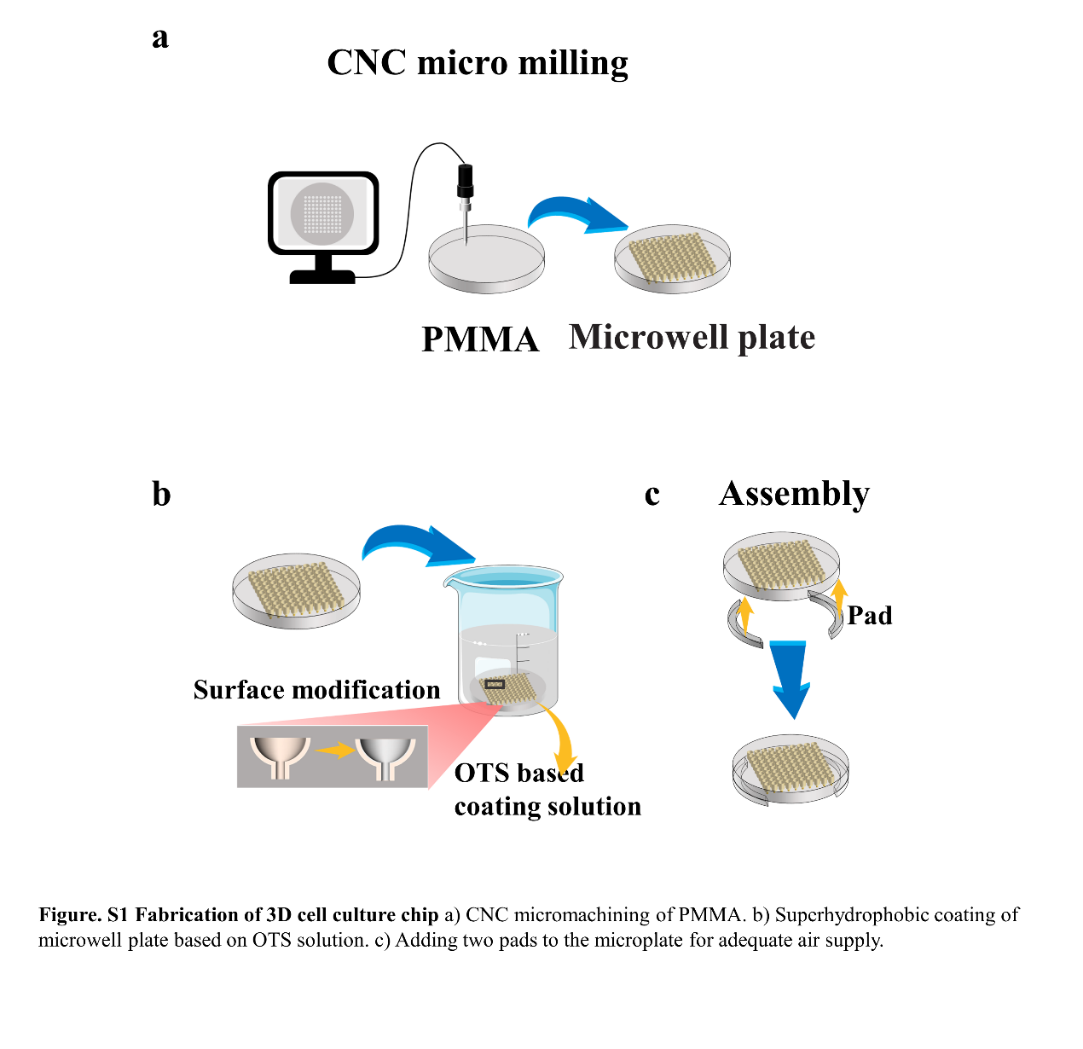

**Figure. S1 Fabrication of 3D cell culture chip** a) CNC micromachining of PMMA. b) Superhydrophobic coating of microwell plate based on OTS solution. c) Adding two pads to the microplate for adequate air supply.

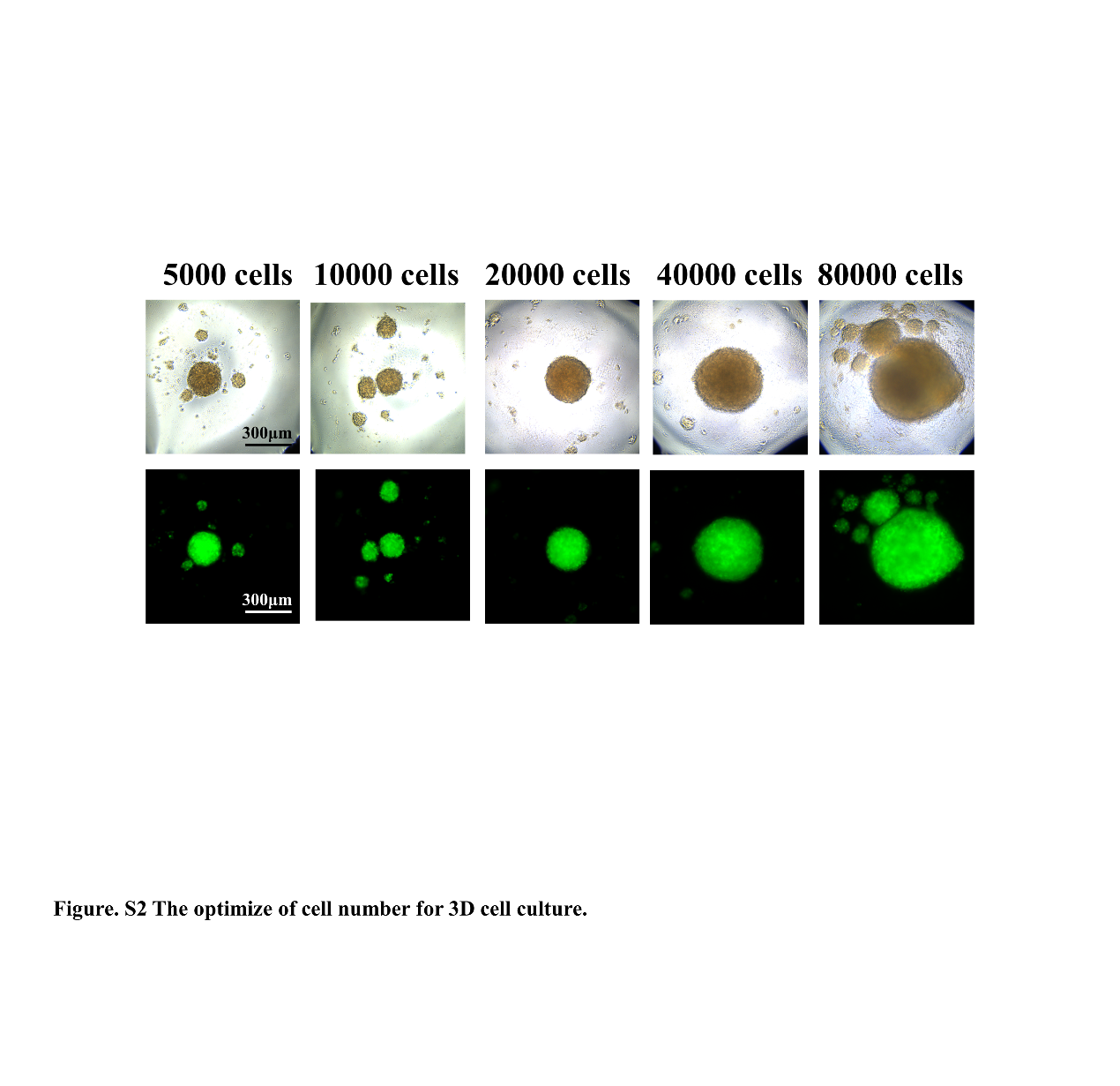

**Figure. S2 The optimize of cell number for 3D cell culture.** The number of the cell was selected from 5000-80000 to identify the reasonable cell quantities for single cell spheroid.

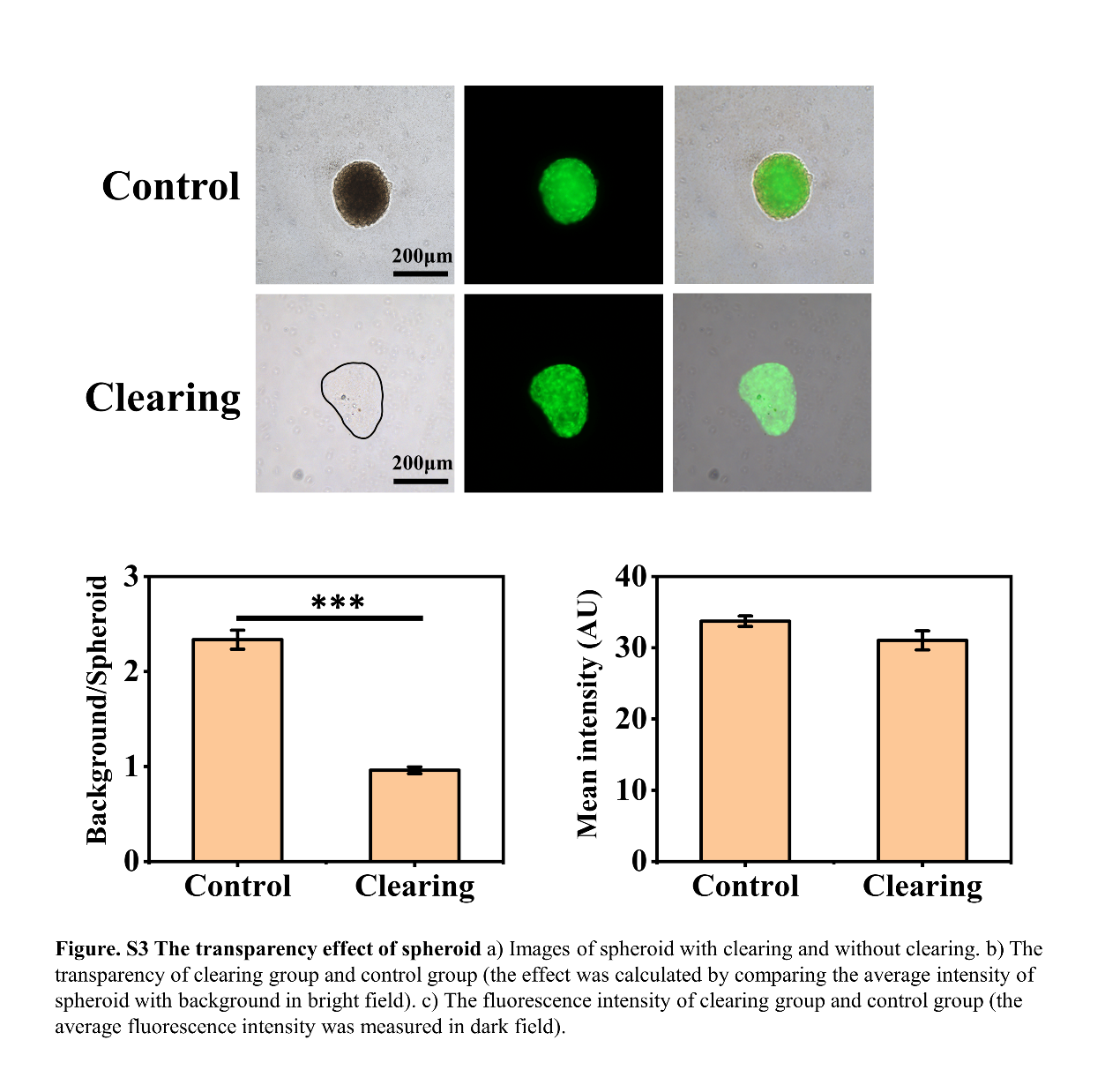

**Figure. S3 The transparency effect of spheroid** a) Images of a spheroid with clearing and without clearing. b) The transparency of the clearing group and control group (the effect was calculated by comparing the average intensity of spheroid with a background in a bright field). c) The fluorescence intensity of the clearing group and control group (the average fluorescence intensity was measured in a dark field).

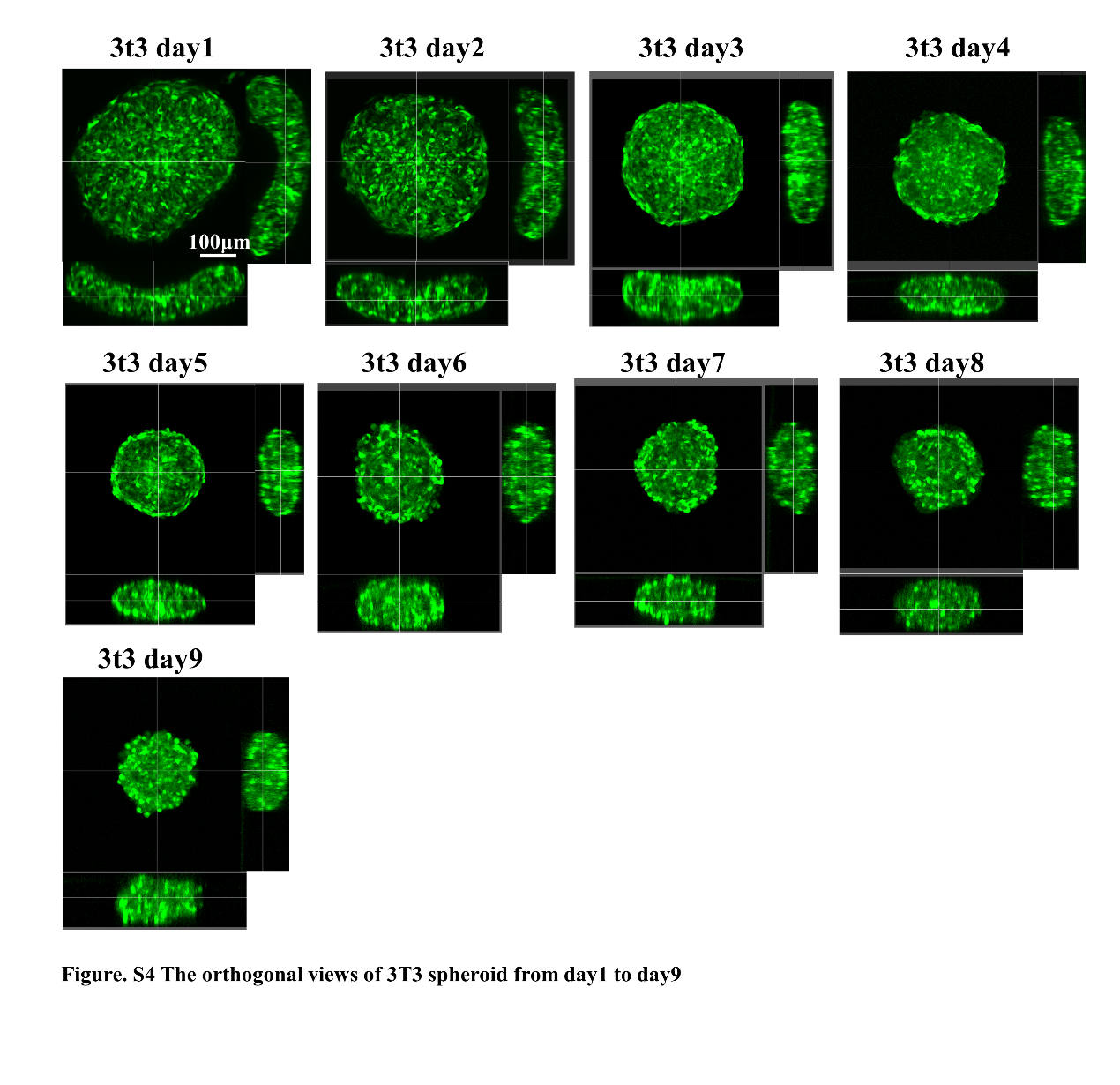

**Figure. S4 The orthogonal views of 3T3 spheroid from day1 to day9**

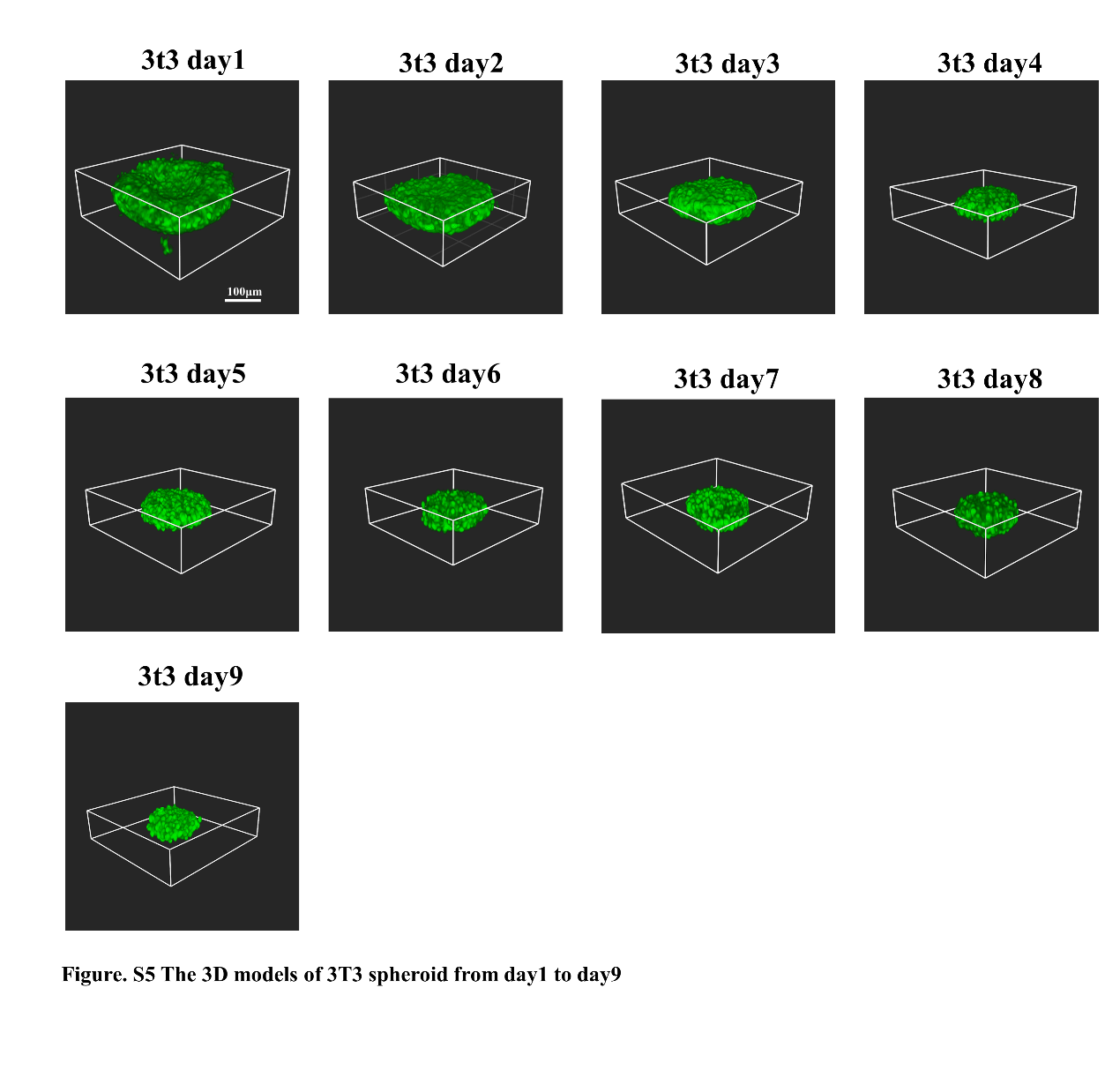

**Figure. S5 The 3D models of 3T3 spheroid from day1 to day9**

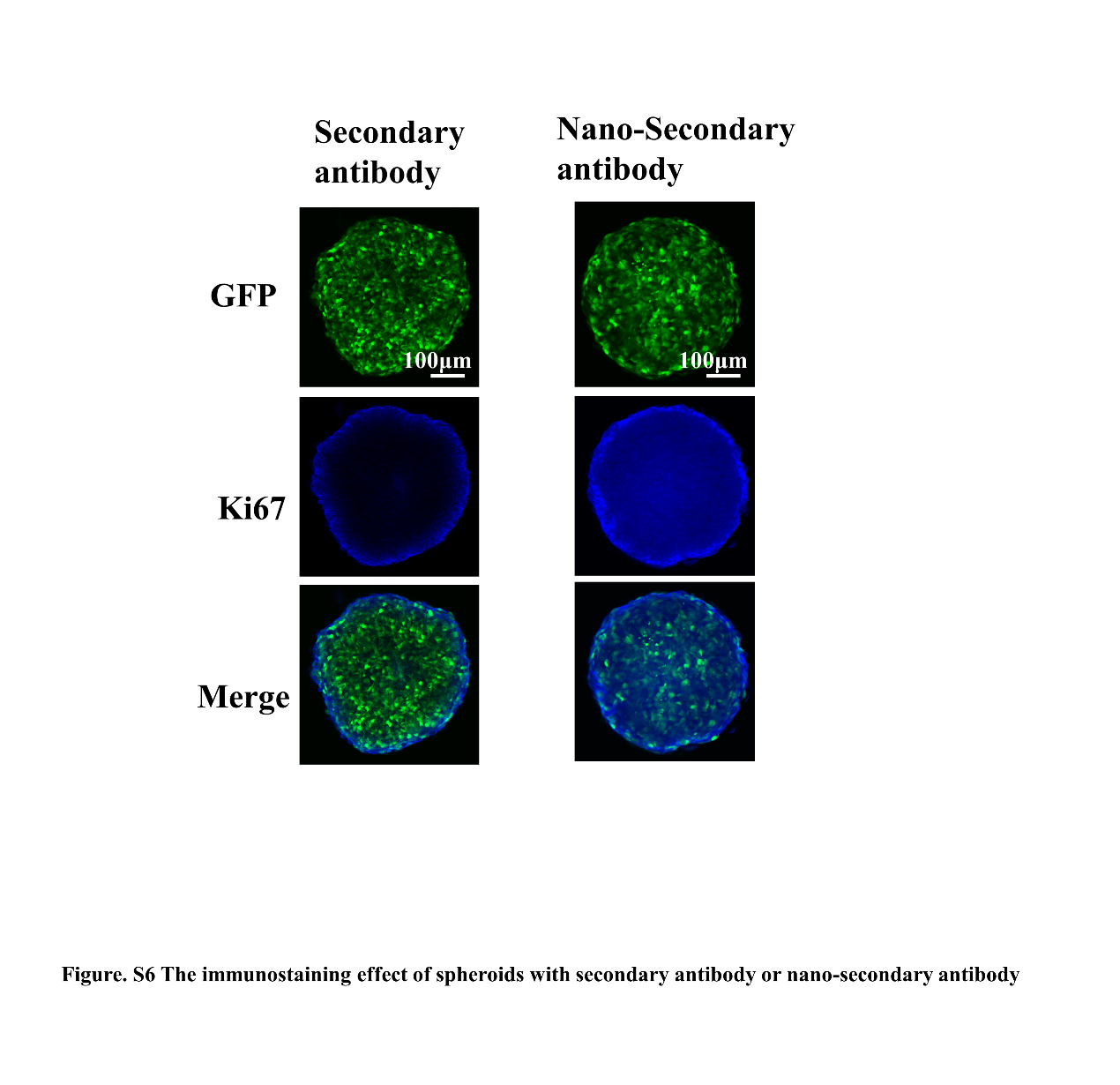

**Figure. S6 The immunostaining effect of spheroids with secondary antibody or nano-secondary antibody.** The spheroids were incubated with normal secondary antibody and nano-secondary antibody (10 times smaller than normal secondary antibody) respectively; it can be found that normal secondary antibody can not penetrate the spheroid while the nano-secondary antibody stained the spheroids evenly.

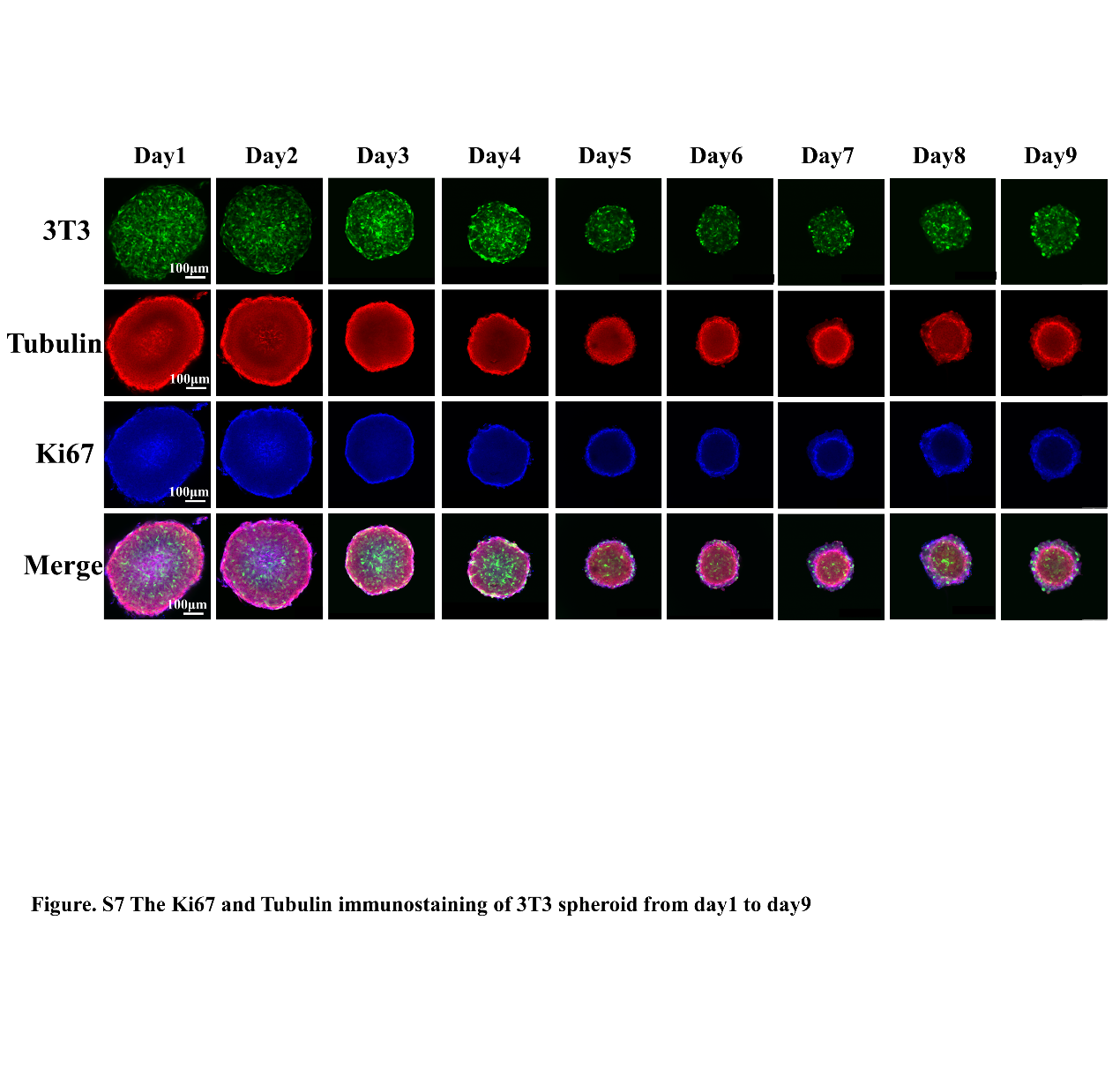

**Figure. S7 The Ki67 and Tubulin immunostaining of 3T3 spheroid from day1 to day9.** The spheroid was treated with primary antibodies (Tubulin and Ki67) from day1 to day7, then incubated with secondary antibodies (red for Tubulin and blue for Ki67).

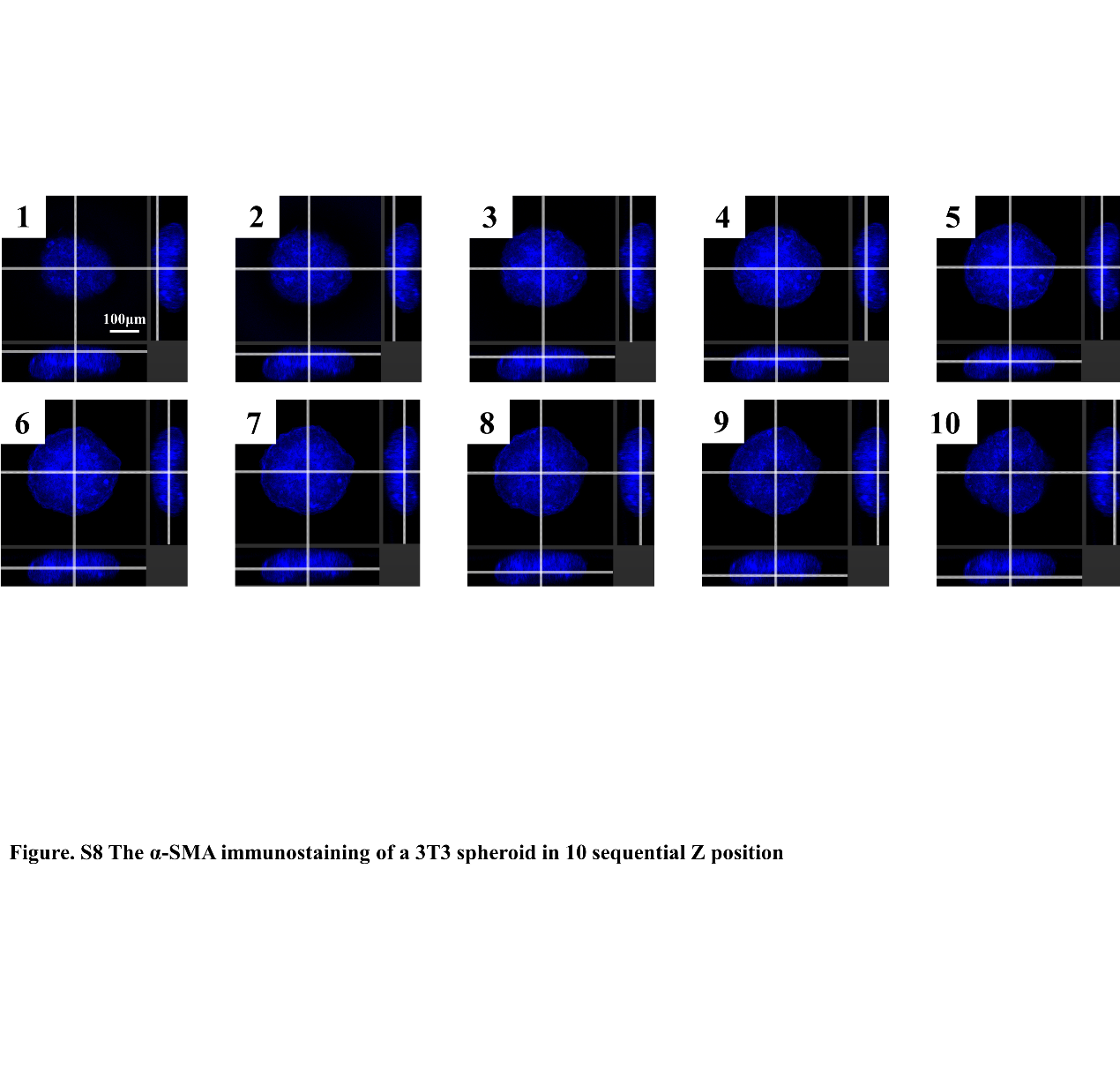

**Figure. S8 The α-SMA immunostaining of a 3T3 spheroid in 10 sequential Z positions.** The spheroid was stained with α-SMA and measured from bottom to top utilizing the z-stack function of confocal microscopy.

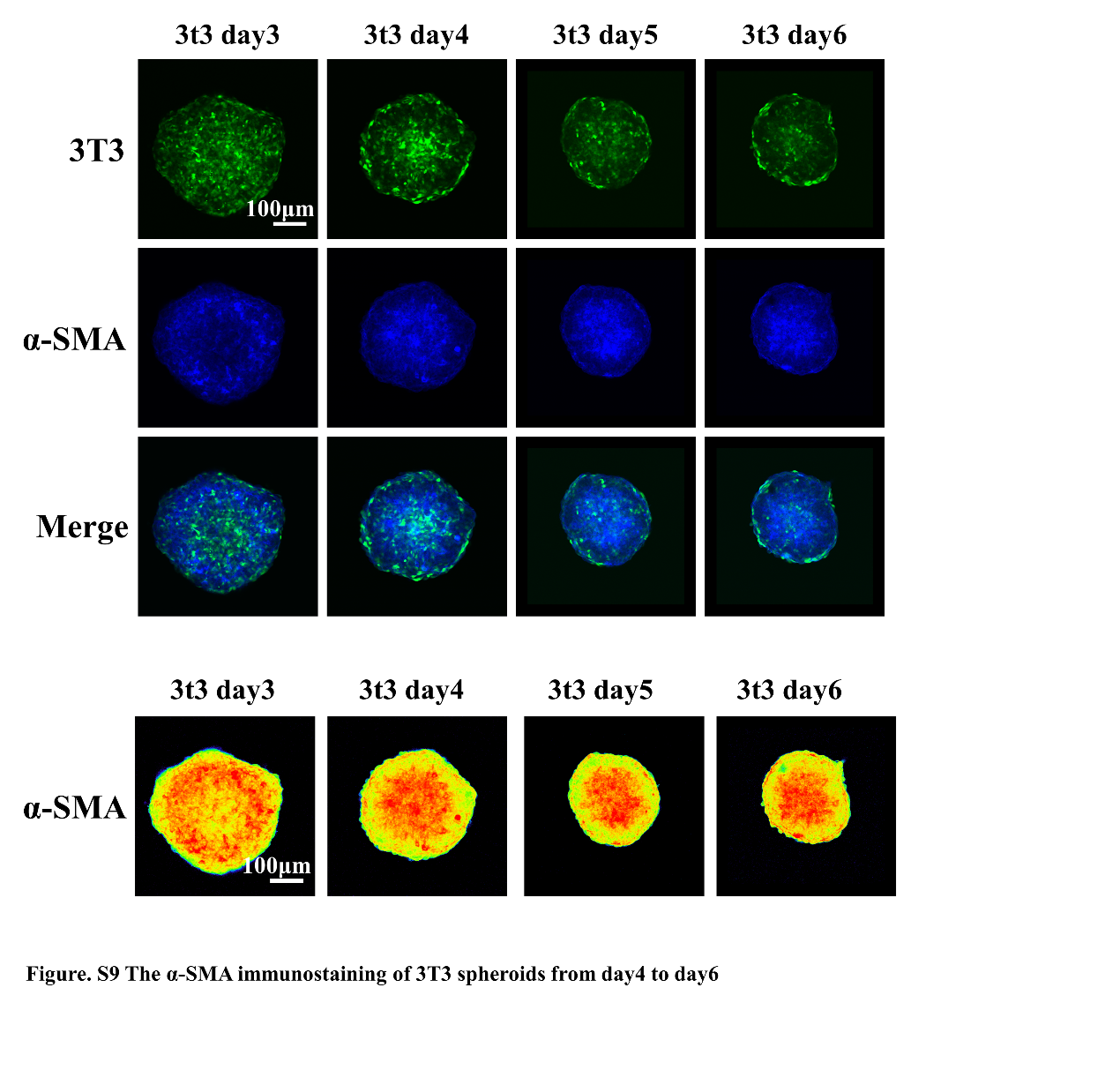

**Figure. S9 The α-SMA immunostaining images of 3T3 spheroids from day3 to day6.** α-SMA was scattered evenly in the 3T3 spheroid in day3; then α-SMA gradually converged to the middle of the spheroid from day4 to day6.

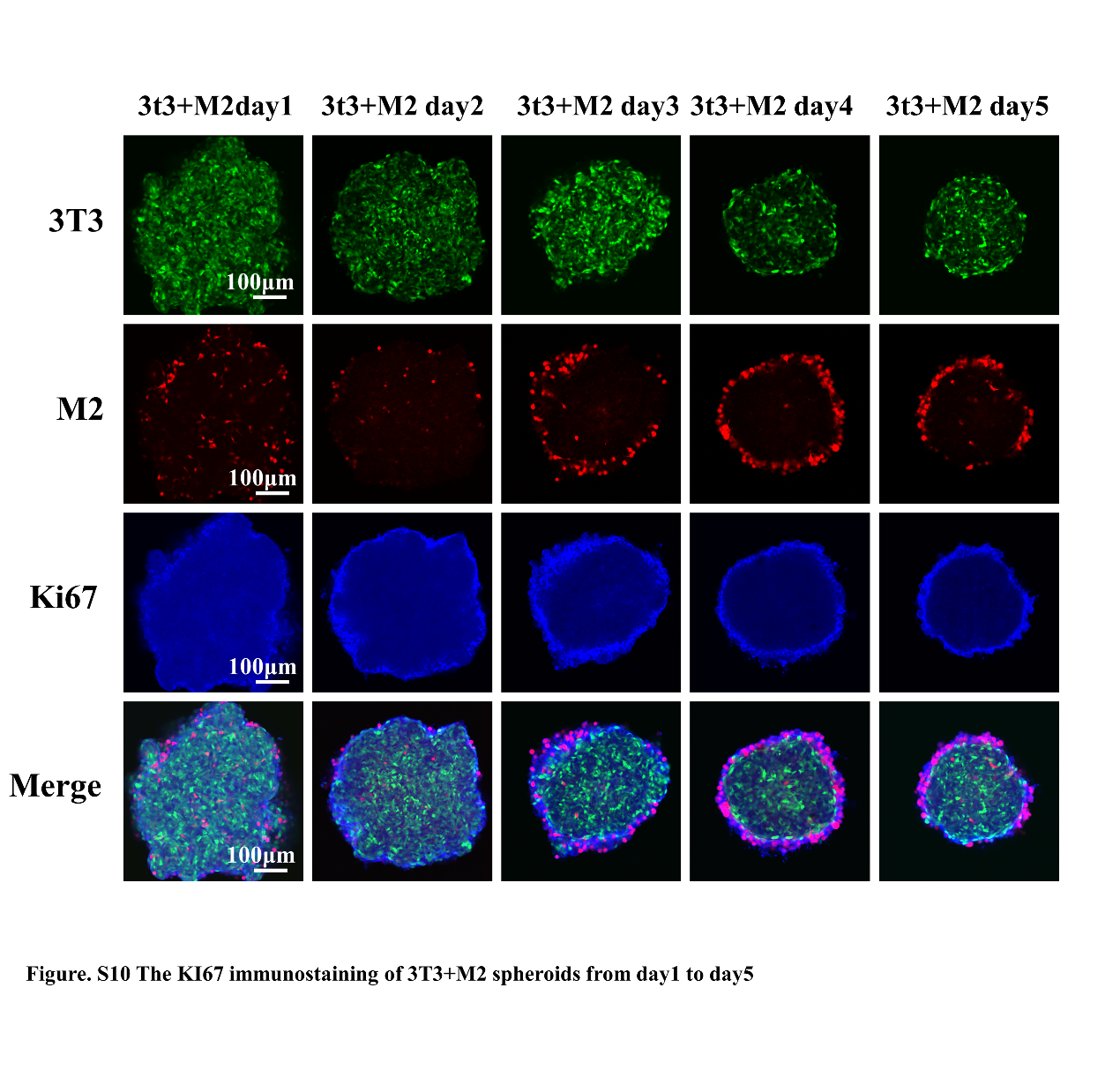

**Figure. S10 The Ki67 immunostaining of 3T3+M2 spheroids from day 1 to day 5.** The 3T3 cells in co-culture spheroid were labeled with GFP and the M2 cells were labeled with RFP. Meanwhile, the spheroid was stained with Ki67. The images suggested that the proliferation of 3T3+M2 spheroids showed no change compared with 3T3 spheroids.
